# Cell free extrachromosomal circular DNA is common in human urine

**DOI:** 10.1101/2021.12.02.471038

**Authors:** Wei Lv, Xiaoguang Pan, Peng Han, Ziyu Wang, Hao Yuan, Weijia Feng, Qingqing Wang, Kunli Qu, Zhe Xu, Yi Li, Tianyu Zheng, Ling Lin, Chengxun Liu, Xuemei Liu, Hanbo Li, Rasmus Henrik Amund Henriksen, Lars Bolund, Lin Lin, Xin Jin, Huanming Yang, Xiuqing Zhang, Birgitte Regenberg, Yonglun Luo

**Affiliations:** College of Life Sciences, University of Chinese Academy of Science, Beijing 100049, China; IBMC-BGI Center, the Cancer Hospital of the University of Chinese Academy of Sciences (Zhejiang Cancer Hospital), Institute of Basic Medicine and Cancer (IBMC), Chinese Academy of Sciences, Hangzhou, Zhejiang 310022, China; Lars Bolund Institute of Regenerative Medicine, Qingdao-Europe Advanced Institute for Life Sciences, BGI-Qingdao, Qingdao 266555, China; BGI-Shenzhen, Shenzhen 518083, China; Department of Biomedicine, Aarhus University, Aarhus 8000, Denmark; Steno Diabetes Center Aarhus, Aarhus University Hospital, Aarhus 8200, Denmark; Department of Biology, University of Copenhagen, Copenhagen 2200, Denmark; Beijing Institutes of Life Science, Chinese Academy of Sciences, Beijing 100101, China; Guangdong Provincial Academician Workstation of BGI Synthetic Genomics, BGI-Shenzhen, Shenzhen, 518120, China; School of Basic Medicine, Qingdao University, Qingdao, 266011, Shandong, China

**Keywords:** Cell free DNA, Noninvasive biomarker, Next generation sequencing, Extrachromosomal circular DNA, DNA circles

## Abstract

Cell free extrachromosomal circular DNA (eccDNA) is evolving as a potential biomarker in liquid biopsies for disease diagnosis. In this study, an optimized next generation sequencing-based Circle-Seq method was developed to investigate urinary cell free eccDNA (ucf-eccDNA) from 28 adult healthy volunteers (mean age = 28, 19 males/ 9 females). The genomic distributions and sequence compositions of ucf-eccDNAs were comprehensively characterized. Approximately 1.2 million unique ucf-eccDNAs are identified, covering 14.9% of the human genome. Comprehensive characterization of ucf-eccDNAs show that ucf-eccDNAs contain higher GC content than flanking genomic regions. Most eccDNAs are less than 1000 bp and present four pronounced peaks at 203, 361, 550 and 728 bp, indicating the association between eccDNAs and the numbers of intact nucleosomes. Analysis of genomic distribution of ucf-eccDNAs show that eccDNAs are found in all chromosomes but enriched in chromosomes i.e. chr.17, 19 and 20 with high density of protein-codding genes, CpG islands, SINE and simple repeat elements. Lastly, analysis of sequence motif signatures at eccDNA junction sites reveal that direct repeats (DRs) are commonly found, indicating a potential role of DRs in eccDNA biogenesis. This work underscores the deep sequencing analysis of ucf-eccDNAs and provides a valuable reference resource for exploring potential applications of ucf-eccDNA as diagnostic biomarkers of urogenital disorders in the future.

**Significance Statement:** Extrachromosomal circular DNA (eccDNA) is an important genetic element and a biomarker for disease diagnosis and treatment. In this study, we conduct a comprehensive characterization of urinary cell free eccDNA (ucf-eccDNA) in 28 heathy subjects. Over one million ucf-eccDNAs are identified. Ucf-eccDNAs are characterized as high GC content. The size of most ucf-eccDNAs is less than 1000 bp and enriched in four peaks resembling the size of single, double, triple, and quadruple nucleosomes. The genomic distribution of ucf-eccDNAs is enriched in generic regions, protein-coding genes, Alu, CpG islands, SINE and simple repeats. Sequence motif analysis of ucf-eccDNA junctions identified simple direct repeats (DRs) commonly presented in most eccDNAs, suggesting potential roles of DRs in eccDNA biogenesis.

## Introduction

Over the past decades, considerable interest has been focused on the clinical application of circulating cell free deoxyribonucleic acid (cfDNA)(1–3). cfDNA molecules are extracellular DNA fragments originated from illness-specific cell death and also normal cell turnover(4). The cfDNA can be detected in a variety of human body fluids, such as urine, saliva, cerebrospinal fluid and blood(5). When cell apoptosis/death occurs, cfDNA is released into the circulatory system and a fraction of them can pass through the glomerular filtration barrier into the urine (also known as trans-renal DNA)(6). Meanwhile, cells lining the urinary tract can also secrete cfDNA into urine directly(7). Since urinary cfDNA derives from both the circulatory system and the urinary tract, it reflects the systemic status of individuals. Previous studies on the fragmentation patterns, copy number aberration, mutation status, methylation, integrity and concentrations of urinary cfDNA revealed that urinary cfDNA has emerged as an informative biomarker for prenatal screening, organ transplantation monitoring and infectious/ urologic malignant disease diagnosis(3, 8–13). However, these studies are all limited to cell free linear DNA in urines.

Extrachromosomal circular DNA (eccDNA) refers to a group of double/single strand circular DNAs that are independent of nuclear chromosomes, but homologous to chromosome DNA(14). To some degree, this special type of DNA molecules reflects the genomic plasticity and stability(15). The eccDNA is commonly presented in all cell types and has been found in many organisms from animals, plants, to microorganisms(16–21). Recent studies have reported that eccDNA is present in murine and human plasma under both cancerous and healthy conditions(22–24). Owing to the covalently closed circular structure of eccDNA, plasma eccDNA is considered more resistant to exonucleases and more stable compared to linear cfDNA. Therefore, plasma eccDNA is expected and has been explored as a novel biomarker for liquid biopsy(23–26).

Urine is an attractive clinical source for early disease detection, especially in urogenital disorders(3, 13). However, the eccDNA profiles in human urine remain unclear, even in healthy individuals. We hypothesized that cell free eccDNA also exits in human urine, because 1) DNA molecules with molecular weights less than 70 kDa (equal to approximately 1.9 kb) have been shown to be freely filtered at the glomerulus and 2) under both pathophysiological and physiological conditions, tissues are known to release eccDNA to the external environment(23, 27). Thus, we modified the Circle-Seq method (14, 18) and conducted a systematic characterization of urinary circular DNA from 28 adult healthy volunteers. To our knowledge, this work, for the first time, systematically characterize the features of cell free eccDNA in human urine and provided comprehensive baseline information for future studies.

## Results

### A modified method for purifying ucf-eccDNAs

To decipher the characteristics of cell free eccDNA in human urine (ucf-eccDNA), we modified the Circle-Seq method which was developed previously for enriching eccDNAs from yeast and human somatic tissues (see methods)(14, 18). Briefly, we isolated cell free DNA (cfDNA) from urine sample using a magnetic beads-based DNA purification approach (see methods). Then, as a positive control of circular DNA, plasmids mix control were added to the total urinary cfDNA before exonuclease digestion. Linear DNAs were then removed from the cfDNA by using Plasmid Safe ATP-dependent exonuclease treatment. The enriched circular DNAs were subjected to rolling circle amplification (RCA) using a highly processive phi29 DNA-polymerase. *MssI* and *NotI* restriction enzymes digestion, which cleaves the spike-in plasmids, showed that the spike-in plasmids control was amplified greatly by our approach (**Figure S1**). In addition to the spike-in plasmids, strong DNA staining signal (smear) was also detected for urine samples but not plasmids control, indicating that eccDNAs are present in the urine samples (**Figure S1**). Based on the optimized protocol of eccDNA purification from urine samples, as depicted in **Figure 1**, we isolated urinary cell free eccDNAs (ucf-eccDNAs) from 28 adult healthy volunteers (mean age 28.9 years (range 22 to 39 years); 19 males/ 9 females). The RCA-enriched ucf-eccDNAs were deep-sequenced with MGI 150-bp pair end (PE) read sequencing. The Circle-Map software, a highly sensitive circular DNA detection method(28), was used to identify ucf-eccDNAs based on split read and discordant read pairs (ref. genome hg38).

**Figure 1.**
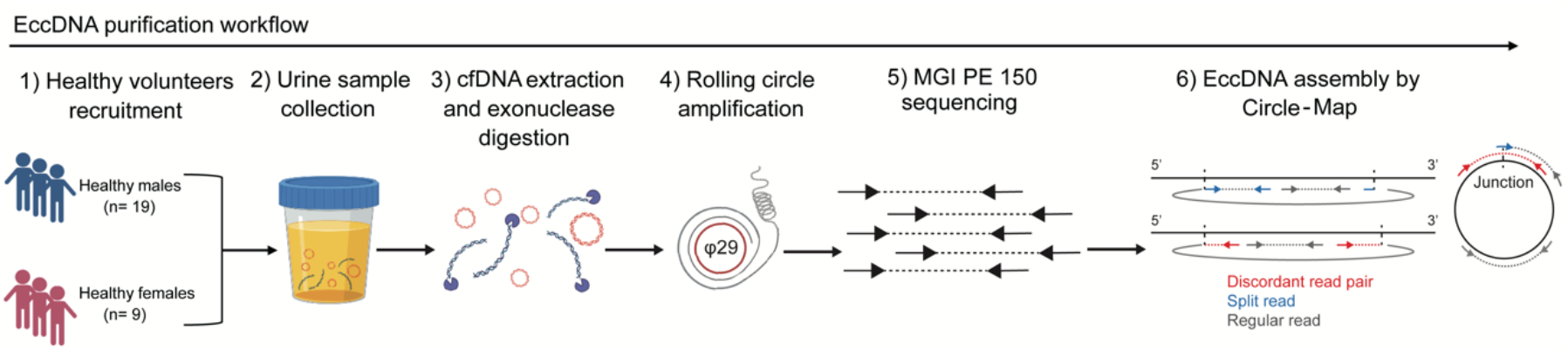
Workflow of cell free eccDNA purification and identification from urine samples. Total cell free DNAs consisting of both circular- and linear-DNAs were isolated from the urine samples. To purify circular DNAs, linear DNA molecules were removed by exonuclease digestion. The enriched circular DNAs were then amplified by rolling circle amplification. The amplified products were finally subjected to library construction and sequencing. Circle-Map software was used to detect eccDNAs from sequencing data.

In total, we detected 1,211,827 unique ucf-eccDNAs from all urine samples and the number of eccDNAs varied appreciably among each urine sample (male, average: 50,948, range 1471-190,738; female, average: 27,088, range 5959-74,965) (**Table S1**). Since the number of detected eccDNAs is affected by the sequencing depth, we normalized it by calculating the numbers of eccDNA per million mapped reads. There was no significant difference between the normalized number of eccDNAs between male and female (p = 0.29) (**Figure S2**). Thus, male and female ucf-eccDNAs were merged for most downstream analyses unless specified. Most (64.05%) ucf-eccDNAs were mapped to genic regions and 50.14% of ucf-eccDNAs were mapped to protein coding genes. The number of eccDNAs also varies greatly between different protein coding genes. Further analysis showed that there was a significantly positive correlation between the length of protein coding genes and the number of eccDNAs (Pearson R = 0.88) (**Figure S3**). Because genic regions and protein-coding genes are better annotated than other DNA regions, to ensure that this was not caused by mapping and annotation issues, we generated 28 random datasets with ~1.2 million *in silico* eccDNAs (size from 150 bp to 850 bp). Unlike the ucf-eccDNAs, only 56.42% and 41.61% of the randomly generated *in silico* eccDNAs were mapped to genic regions and proteincoding genes, suggesting that ucf-eccDNAs are more frequently derived from generic regions. Since all the ucf-eccDNAs are from healthy subjects in this study, we focus on charactering the genomic and sequence features of the eccDNAs, thus providing a valuable reference resource for future investigations.

### Guanine-cytosine (GC) content and size distributions of ucf-eccDNAs

We first analyzed the GC content of ucf-eccDNAs as it is well known that GC content is associated with DNA stability, structure, evolution, and functions(29). The overall GC content of the human genome is about 45% and previous studies have reported that eccDNA molecules are more likely originated from genomic regions with high GC content(21, 30). The GC content of ucf-eccDNAs was significantly higher than that of their downstream and upstream regions of equivalent length and the *in silico* eccDNAs (**Figure 2A, B**), however this high GC preference was not seen in the randomly generated eccDNA dataset. The GC content distribution of ucf-eccDNAs from each individual sample was displayed in **Figure S4**, which shows consistently higher GC contents in the ucf-eccDNA than the flanking sequences.

**Figure 2.**
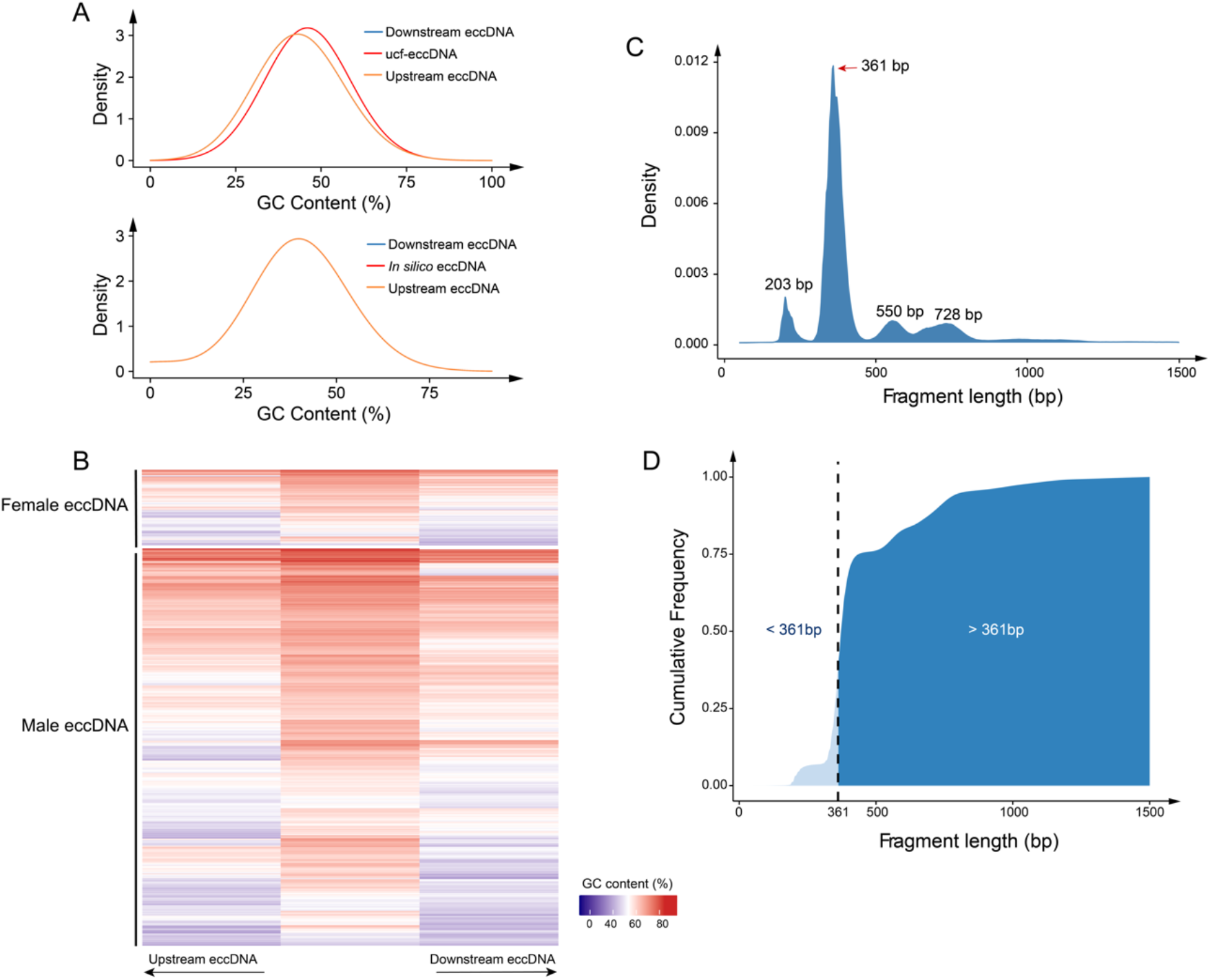
GC content and length distributions of urinary cell free eccDNAs. (A) GC content distribution of urinary cell free/ in silico eccDNAs and their downstream and upstream regions of equivalent length. (B) Heatmap showing GC content of randomly selected 4000 urinary cell free eccDNAs and their downstream and upstream regions of equivalent length. (C) Length distribution of urinary cell free eccDNAs. (D) Cumulative frequency plot of urinary cell free eccDNAs. Ucf-eccDNA: urinary cell free eccDNA.

We next analyzed the size distribution of ucf-eccDNAs. Most ucf-eccDNAs (99.9%) were smaller than 1000 bp. Interestingly, ucf-eccDNAs are enriched in four characteristics peaks positioned at 203, 361, 550 and 728 bp with a periodicity of ~160-200 bp. Among them, the 361-bp peak was the most pronounced, being about more than ten-fold larger than the other three peaks (**Figure 2C and 2D, Figure S5**). Such size distribution pattern of ucf-eccDNAs was highly conserved across most individual urine samples (**Figure S6**), which is reminiscent of the size of single, double, triple, and quadruple nucleosomes and consistent with previous observations of eccDNAs from plasma and tissues (16, 21–23, 31).

### Effects of Genomic context in cis on ucf-eccDNAs

We next examined the genomic context of ucf-eccDNAs. The ucf-eccDNAs were widely distributed across the human genome (**Figure 3 and S7**). We then combined and merged eccDNA sequences and found that these molecules contained about 14.9% (462.1 Mb) of the total human genomic information, supporting that a considerable proportion of the human genome can form eccDNAs. Meanwhile, we observed that the number of eccDNAs per Mb on each chromosome was uneven. In specific, chromosomes 19 that is gene-rich hosted more eccDNAs than other chromosomes, whereas gene-poor chromosome 13 hosted less (**Figure 4A and 4B**). Consistently, there was positive correlation between the frequency of eccDNA and the number of protein coding genes per Mb (Male, p = 1.79E-3, Pearson R = 0.6; Female, p = 7.99E-4, Pearson R = 0.65), which was consistent with our previous reports(16) (**Figure 4C**). Compared with other chromosomes, chromosome 19 is enriched with *Alu* elements. We therefore hypothesized that *Alu* elements could influence DNA circularization. Our analysis showed that there is a positive correlation between the frequency of eccDNA and the number of *Alu* elements per Mb (Male, p = 4.66E-4, R = 0.66; Female, p = 8.91E-5, R = 0.73) (**Figure 4D**). Most importantly, in the in silico generated random eccDNA datasets, there was no significant correlation between the frequency of eccDNA and the number of protein coding genes per Mb (p = 0.11, R = 0.33) or the number of *Alu* elements per Mb (p = 0.14, R = 0.31).

**Figure 3.**
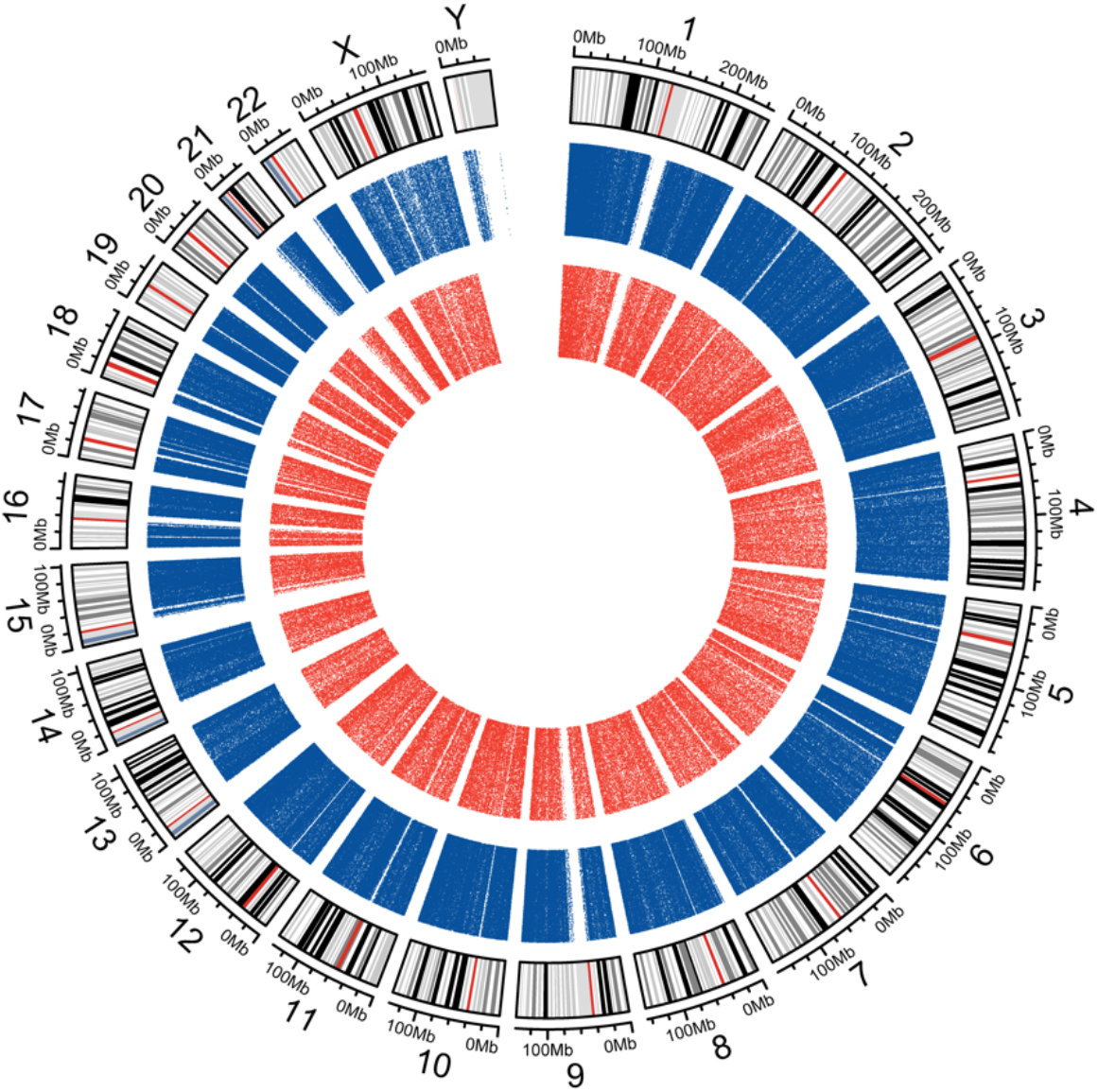
Circos plot presenting the genome-wide distribution of urinary cell free eccDNAs. From inside to outside, the red and blue dots represent female-derived urinary cell free eccDNAs (pooled data from 9 cases) and male-derived urinary cell free eccDNAs (pooled data from 19 cases) respectively.

**Figure 4.**
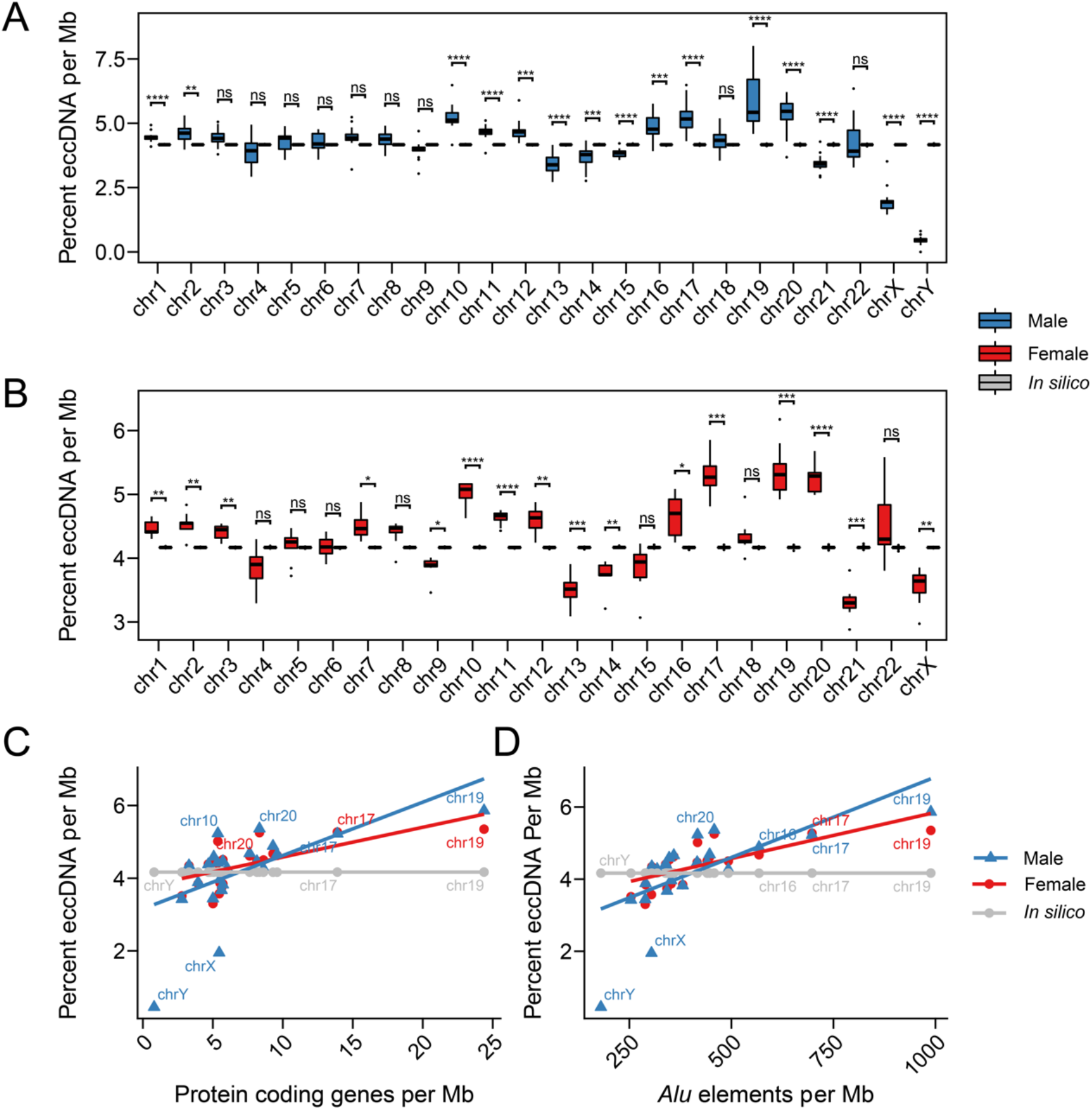
EccDNA frequency correlates with chromosome and the density of protein coding genes, and *Alu* elements. (A, B) The percent of the number of ucf-eccDNAs per Mb in different chromosomes in male (A) and female (B). (C, D) Significant positive correlation between the frequency of eccDNA with the number of (C) protein coding genes per Mb and (D) the number of *Alu* elements per Mb (chromosome X, Y, 10,16, 17, 19 and 20 are marked). ns, not significant; *, **, ***, and **** denote significant p value less than 0.05, 0.01, 0.001, and 0.001 respectively.

### Genomic distribution of ucf-eccDNAs

We further explored the genomic distribution of ucf-eccDNAs from the human genome. To do this, we mapped the ucf-eccDNAs junctions to seven major classes of genomic elements: untranslated regions (UTR), CpG islands (CpG), exons, introns, and 2 kb regions upstream/downstream of genes. Since the length of these genomic elements varied widely, using the absolute number of eccDNA in each type of genomic elements for statistics does not truly reflect its generation preference. Therefore, we used the “observed/expected ratio of genomic elements” to statistically evaluate the enrichment or depletion of ucf-eccDNAs. The “observed/ expected ratio of genomic elements” was calculated as the number of eccDNA falling in a certain type of genomic element divided by the percentage of the length of that genomic element over the length of all these seven types of elements. Surprisingly, we found that the distribution of eccDNAs on the whole genome was not stochastically distributed. Consistent with the above analysis, certain genomic regions are more vulnerable to ucf-eccDNA formation. In particular, compared with *in silico* randomly generated eccDNAs, ucf-eccDNAs were primarily enriched in CpG islands and 5’UTR, whilst significantly (p < 0.001) less were found in the intronic regions (**Figure 5A**).

**Figure 5.**
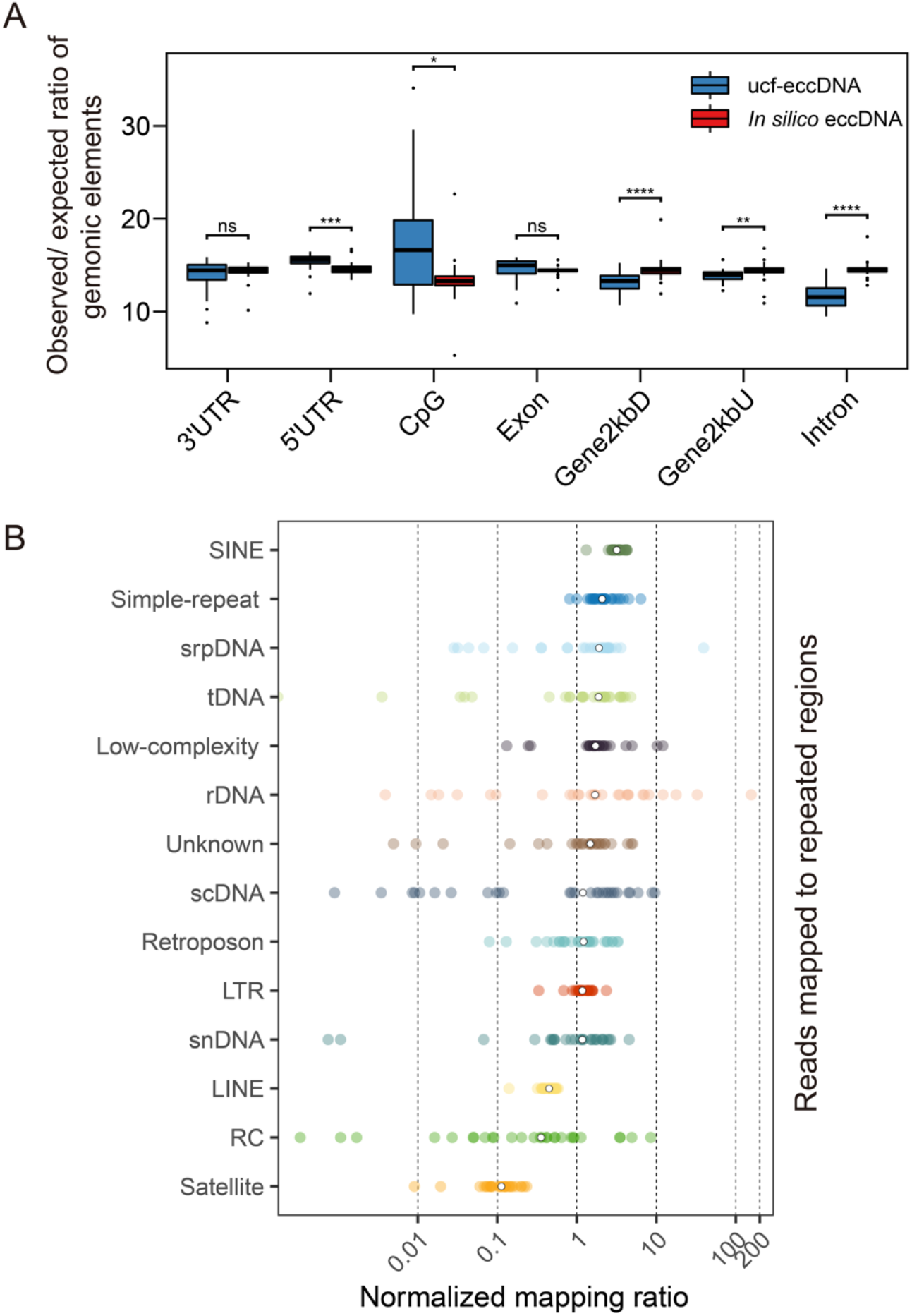
Genomic distributions of urinary cell free eccDNAs. (A) Distribution of urinary cell free eccDNAs in indicated genomic elements. The “observed/ expected ratio of genomic elements” is the number of eccDNA falling in a certain type of genomic element divided by the percentage of the length of that genomic element over the length of all these seven types of elements. (B) Normalized mapping ratio of eccDNA reads in specific repetitive elements (median, white dot). UTR, untranslated region; Gene2kbD, 2kb downstream of genes; Gene2kbU, 2kb upstream of genes; SINE, short interspersed nuclear element; srpDNA, signal recognition particle DNA repeats; rDNA, ribosomal DNA repeats; tDNA, transport DNA repeats; scDNA, small conditional DNA repeats; snDNA, small nuclear DNA repeats; LTR, long terminal repeat; LINE, long interspersed nuclear element; RC, rolling circle repeats. Ucf-eccDNA: urinary cell free eccDNA. ns, not significant; *, **, ***, and **** denote significant p value less than 0.05, 0.01, 0.001, and 0.001 respectively.

Repetitive sequences account for 52.5% of the human genome. In human cells and tissues, repetitive sequences have been reported more prone to generate eccDNAs(16, 19, 32). Therefore, we extracted ucf-eccDNA reads from repetitive sequences in an attempt to gain more insights into the correlation between ucf-eccDNAs and repetitive sequences. The average proportion of ucf-eccDNA reads that aligned to repetitive elements was 70.7%, corroborating previous finding and support the over presentation of ucf-eccDNAs in repetitive sequences. Moreover, we noticed that the enrichment of each repetitive class on ucf-eccDNA was uneven. Specifically, elements encoding short interspersed nuclear elements (SINEs) (3.2-fold, median), simple repeats (2.1-fold, median), signal recognition particle DNA repeats (srpDNA) (1.9-fold, median), tDNA repeats (1.8-fold, median), low complexity repeat (1.7-fold, median) and rDNA repeats (1.7-fold, median) were clearly overrepresented, whereas satellites (0.1-fold, median), rolling circle repeats (RC) (0.4-fold, median), and long interspersed nuclear elements (LINEs) (0.4-fold, median) were less than expected (**Figure 5B**). Collectively, our results suggest that certain genomic contexts such as CpG islands and generic regions are more vulnerable to the generation of eccDNAs.

### Simple Direct repeats (DRs) are commonly found at ucf-eccDNAs junctions

Finally, we sought to characterize the sequence motifs of ucf-eccDNAs as the generation of eccDNA in cells are associated with the endogenous DNA repair machinery. Previously, we demonstrated repair of DNA double strand breaks introduced by CRISPR can lead to the generation of eccDNAs in cells(33). We focus on eccDNA start/end sites (junctions), which was the upstream/downstream edge in the genome domains of eccDNA origin, might related to circularization of certain genome domains. Therefore, we scrutinized the DNA sequences from 10 bp flanking at eccDNA start and end sites in order to investigate the mechanism underlying the formation of ucf-eccDNAs. Notably, we observed 66.36% of the total eccDNA molecules possessed 4- to 18-bp direct repeats (DR) respectively near eccDNA junction sites. As one example, [*RBFOX1*^circle 7,712,600-7,712,978 bp^] with fragment size of 378 bp had 5 bp DR near the eccDNA junction site (**Figure 6A**). Moreover, we also found a pair of trinucleotide fragments (dual-direct-repeat pattern) with 4-bp “spacers” in between was flanking the eccDNA start and end sites, which was similar to the motif signature of plasma eccDNA (**Figure 6B**)(22). We then divided eccDNA molecules into bins by their peak sizes and found that this nucleotide motif pattern was also conserved regardless of fragment size (**Figure S8**), suggesting a potential role of DRs in the biogenesis of eccDNAs.

**Figure 6.**
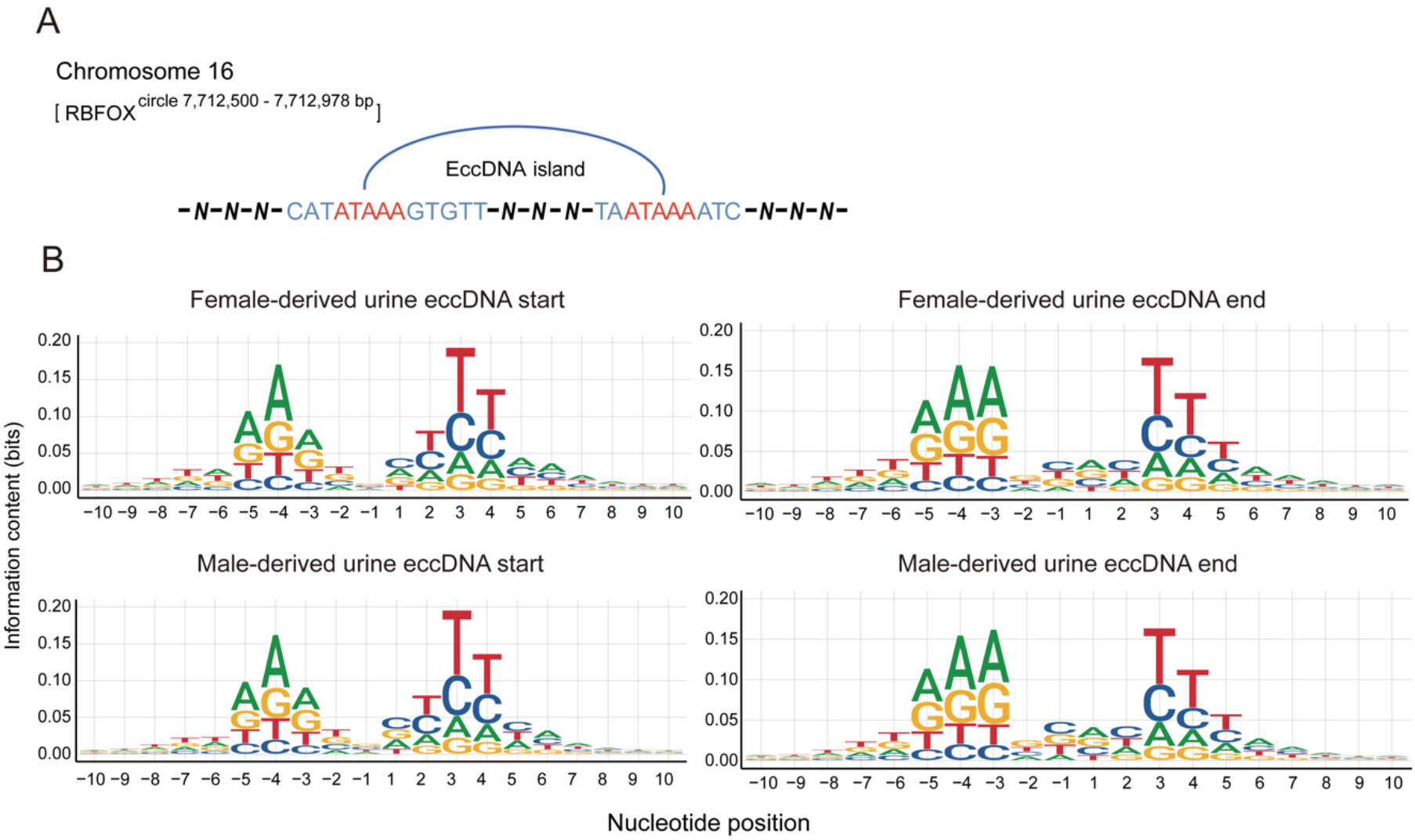
Characterization of urinary cell free eccDNAs fragment junction sites. (A) As one example, [*RBFOX1*^circle 7,712,600-7,712,978 bp^] with 5 bp direct repeats (DR) flanking its start and end sites. (B) The nucleotide frequencies surrounding the start and end sites of male-derived urinary cell free eccDNAs (pooled data from 19 cases) and female-derived urinary cell free eccDNAs (pooled data from 9 cases).

## Discussion

The eccDNA is commonly found in plasma and somatic tissues(16, 21–23). However, its presence in human urine is still largely uncertain. In this study, we demonstrated for the first time that cell free eccDNAs are commonly found in urine. These ucf-eccDNAs are likely derived from the urinary tract, as well as plasma eccDNAs which may cross the filtration barrier into Bowman’s capsule. It also implies a possible manner for the clearance of circulating circular DNA. Though several studies have reported the existence and characteristics of plasma eccDNA, it has been unclear how eccDNA is cleared from the plasma(22–25). Our results suggest that filtration of eccDNA in plasma to the urinary system might be the route for eccDNA to be eliminated from the body.

We identified over one million of ucf-eccDNAs and these eccDNA molecules are widely distributed across the human genome. The combined eccDNA lengths covered 14.9% of the human genome, suggesting a considerable proportion of the genomic regions can form eccDNA. In human somatic tissues, rDNA repeats, tDNA repeats are more likely generate circles than any other repetitive elements, and satellites on eccDNAs was also overrepresented(16, 19). However, in human urine, satellites on eccDNAs were clearly less than expected (0.1-fold, median), and SINEs seem to give rise to circles more frequently than rDNA repeats and tDNA repeats, suggesting that eccDNA formation exhibited genomic context-specific vulnerability.

It has been reported that urine-derived cell free linear DNA is highly degraded, with its modal fragment length ranging from 80-110 bp(3, 34). Here we found the eccDNA number among each urine sample varied widely. We therefore speculated that cell free eccDNAs are also degraded to some extent in the urine, although circular DNA is more stable than linear DNA. Further study is needed to figure out the degradation pattern of urine cell free eccDNAs. Moreover, ucf-eccDNAs were significantly longer than its linear counterpart. The fragment length distribution of ucf-eccDNAs presented four appreciable peaks clusters located at 203, 361, 550 and 728 bp (**Figure 2C**). Such length profile is quite similar to the length distribution of plasma eccDNAs(21, 22). The fragment patterns of urinary cfDNA were previously proven to be associated with nucleosome degradation(3). The nucleosome consists of two linkers (~40 bp) and a core region (~146 bp). We noticed that ucf-eccDNAs showed a 160-200 bp periodicity, suggesting that the eccDNA pattern may derive from nucleosomes.

The underlying mechanism regulating eccDNA generation remains unknown and likely involves various mechanistic models. Some models suggested that microhomology directed repair of double-stranded DNA (e.g., microhomology mediated end joining and homologous recombination) may mediate eccDNA formation because a large portion of circles had various repeated sequences (4-18 bp) at their junction sites(15, 16, 18). However, some eccDNA molecules did not contain any repeats and thus it is possible that other pathways regulated circularization. EccDNA production was strongly associated with DNA replication, and neither ours nor previous studies found that transcriptionally active and GC-rich domains are more prone to generate eccDNA. Moreover, dual direct repeats flanking the junction sites of eccDNA were observed in this study and this motif pattern was quite similar to that in plasma eccDNA of pregnancy. All these features indicated that R-loop formation and replication slippage may also lead to the eccDNA formation (based on single-stranded DNA)(15, 22). Overall, multiple pathways seem to mediate the formation of eccDNA.

In summary, in this work we found that cell free eccDNA is common in human urine and systematically described the features (e.g., genomic distribution, size profiles, GC content, and motif signatures) of these special types of DNA molecules. Compared with urinary cell free linear DNA, eccDNAs had significantly longer length and distinct motif pattern. These interesting features of urine cell free eccDNAs may prompt future studies for biomarker development of multiple urogenital disorders. Collectively, we provide valuable insights into the presence, distribution, and compositions ucf-eccDNAs in healthy subjects. This provides an important reference for the further exploration of ucf-eccDNAs as diagnostic biomarkers of human diseases.

## Materials and Methods

### Healthy volunteers’ recruitment and urine sample collections

The aim of this study was to explore the presence of cell free eccDNA in human urine. Twenty-eight healthy volunteers (mean age 28.9 years (range 22 to 39 years); 19 males/ 9 females) without any urological system disorders were recruited from the Qingdao-Europe Advanced Institute for Life Sciences and signed an informed consent approved by the Institutional Review Board (IRB) of BGI-Shenzhen (BGI-IRB) (**Table S1**). The day before urine sample collections, all volunteers were asked to prohibit from drinking water for 12h. Approximately 30 ml of first morning-voided midstream urine sample added with 0.6 ml of 500mM EDTA was collected between 7am and 8am on each day of donation. The urine samples were stored at 4 degree for further processing. During the sample collection processes, all urine samples from female participants were collected by the participants at their home using a self-collection urine kit. But all samples were delivered to our laboratory within one hour at 4 degree.

### cfDNA isolation from urine samples

To minimize cfDNA degradation, all urine samples were processed within two hours after collection. To obtain the cell free portion and remove cellular matter, 10 ml of urine were spun 16000 x g for ten minutes at 4 degree and the resulting supernatant fluids were passed through a 0.45 μm filter. cfDNA was isolated from 5 ml cell free urine supernatant using a MGIEasy Circulating DNA Extraction Kit (MGI-BGI, China) and eluted in 55 ul RNase-free water. The cfDNA concentration was measured by Qubit dsDNA HS Assay kit (Invitrogen).

### Plasmids

The pAAV-saCas9-Backb (P895; 7447 bp) and BE4-Gam (P1035; 9444 bp) plasmids were purified from Escherichia coli using the TIANGEN mini plasmid purification kit.

### Cell free eccDNA purification and amplification

To remove the linear portions of urinary cfDNA, 40 ul of urinary cfDNA added with *spike-in* plasmids control (~ 100 copies of P895 and ~ 100 copies of P1035) was treated with 25 unites of Plasmid-Safe DNase (Epicenter) at 37°C for 24 hours. The Plasmid-Safe DNase was then inactivated by incubating samples at 70°C for 30 minutes. The digestion products were cleaned up from reaction mixes using 2.0× WAHTS® DNA clean Beads (Vazyme). Approximately 50% (12 μL) of the total volume of resultant circular DNA was subjected to rolling circle amplification (RCA) reaction with incubation at 30°C for 16 hours. The RCA reaction system involved 1 ul Phi29 polymerase (Thermo), 4 ul 10× Phi29 buffer (Thermo), 2 ul exonuclease-resistant random primer (Thermo), 4 ul 2.5 μM dNTP mixture (Takara), 0.8 ul 100mM DTT and 14.2 ul RNase-free water. The phi29-amplified products were recovered by 2.0× WAHTS® DNA clean Beads, eluted in 80 ul RNase-free water and quantified by Qubit 3.0.

### Quality control by agarose gel electrophoresis

Successful amplification was assessed by agarose gel electrophoresis (0.7%). Moreover, 1 ul of RCA products were double-digested with *MssI* and *NotI* restriction enzymes (Thermo) for 30mins at 37°C. The digestion products were also subjected to agarose gel electrophoresis (0.7%).

### Library preparation and sequencing

About 500 ng of amplified DNA were sheared by sonication (Covaris LE220) to generate a median size of 400 bp. Then, 50 ng of fragmented DNA were input for library construction using MGIEasy DNA Library Preparation Kit (MGI-BGI, China). Length distribution and quality of each library were examined by the Bioanalyzer 2100 (Agilent). DNA libraries were sequenced (PE150) on MGISeq-2000 platform (BGI-Qingdao, Qingdao, China).

### eccDNA identification from Circle-Seq data

In order to detect eccDNA from PE150 high throughput sequencing data, we applied the Circle-Map software (https://github.com/iprada/Circle-Map)(28). Sequencing reads were firstly aligned to the human reference genome (hg38 genome download from UCSC) using BWA-MEM. Then, two BAM files were sorted according to reads name and coordinates for the extraction of circular reads, respectively. The above files were lastly used for detect coordinates of each eccDNA. In order to improve the accuracy of eccDNA detection, several filtering steps were conducted. The specific settings are as follows: (1) split reads ≥ 2, (2) discordant reads ≥ 2, (3) Circle score ≥ 50, (3) Coverage increase in the start coordinate ≥ 0.33, (4) Coverage increase in the end coordinate ≥ 0.33, (5) Coverage continuity ≤ 0.5, (6) fold between split reads and discordant reads ≤ 20. The detailed information of each parameter is described by Iñigo et al(28).

### Genomic features acquirement

The genomic features, such as the length of each chromosome, the number of Alu-elements, pseudogenes, non-coding RNAs and coding genes in each chromosome, and the position of each gene on chromosomes, were retrieved from Ensembl database assembly GRCh38.

### Genomic annotation of ucf-eccDNAs

The genomic annotation data was downloaded from the UCSC table browser (https://genome.ucsc.edu/cgi-bin/hgTables/). Seven major classes of genomic elements (3’UTR, 5’UTR, CpG island, exon, Gene2KbD, Gene2KbU and intron) were used to map ucf-eccDNAs with junction site. We used the “observed/ expected ratio of genomic elements” for statistics and the “observed/ expected ratio of genomic elements” was calculated according to the following formula:

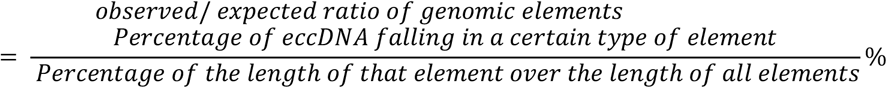

Repetitive sequences analysis was performed as our previously reported(19). Briefly, using BedTools multicov and BedTools group by, we counted the number of reads mapped to the specific repetitive element (RepeatMasker open-4.0.5; http://repeatmasker.org). The normalized mapping ratio was calculated as the percent of reads mapped to the specific repetitive element divided by the percent of the specific repetitive in the nuclear genome.

### Generation of random eccDNA datasets

Based on the weighted average of chromosome length, we generated 28 random datasets of *in silico* eccDNAs across the genome. The number of *in silico* eccDNAs in each dataset was in one-to-one correspondence with the number of ucf-eccDNAs in each urine sample. To better emulate our detected ucf-eccDNAs, the size of *in silico* eccDNAs was range randomly between 150 bp to 850 bp.

### Characterization of ucf-eccDNA fragment junction site

Bedtools-2.29.2 was used to extract 10 bp up- and down-stream DNA sequences of each eccDNA junction site. HomerTools was used to calculate the mean per base mononucleotide frequencies. The motif signature of eccDNA fragment junction site was visualized by R package ggseqlogo-0.1.

### Statistical Analysis

All statistical tests were implemented by R-4.1.1. The difference comparison of two groups was performed by Wilcoxon-test. A Pearson correlation test was subjected to correlation analysis. P < 0.05 indicated statistical significance.

## Acknowledgments

The urine cell free eccDNA project was partially supported by Qingdao-Europe Advanced Institute for Life Sciences. We thank the China National GeneBank for the support of executing the project under the framework of Genome Read and Write. B.R and Y.L are also supported by the CIRCULAR VISION project from the European Union’s Horizon 2020 program and innovation programme under grant agreement No. 899417. We thank Dr. Fred Dubee for his comments and corrections to the manuscript.

## Notes

### Competing Interest Statement

The authors have declared no competing interest.

