## Supplementary figures and Tables for "Cell free extrachromosomal circular DNA is common in human urine"

**This PDF file includes:**

Figures S1 to S8
Tables S1

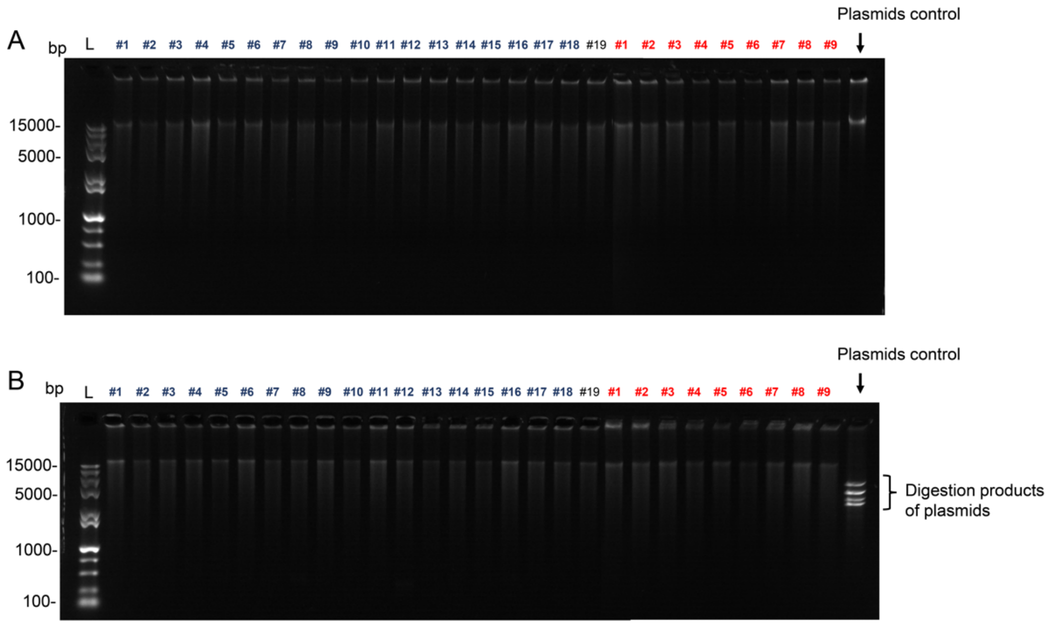


**Fig. S1.** Quality control of RCA products by agarose gel electrophoresis (0.7%). (A) Gel image of RCA products. (B) Gel image of the double-digested RCA products with MssI and NotI restriction enzymes. Urine samples from male and female were marked with blue and red respectively.

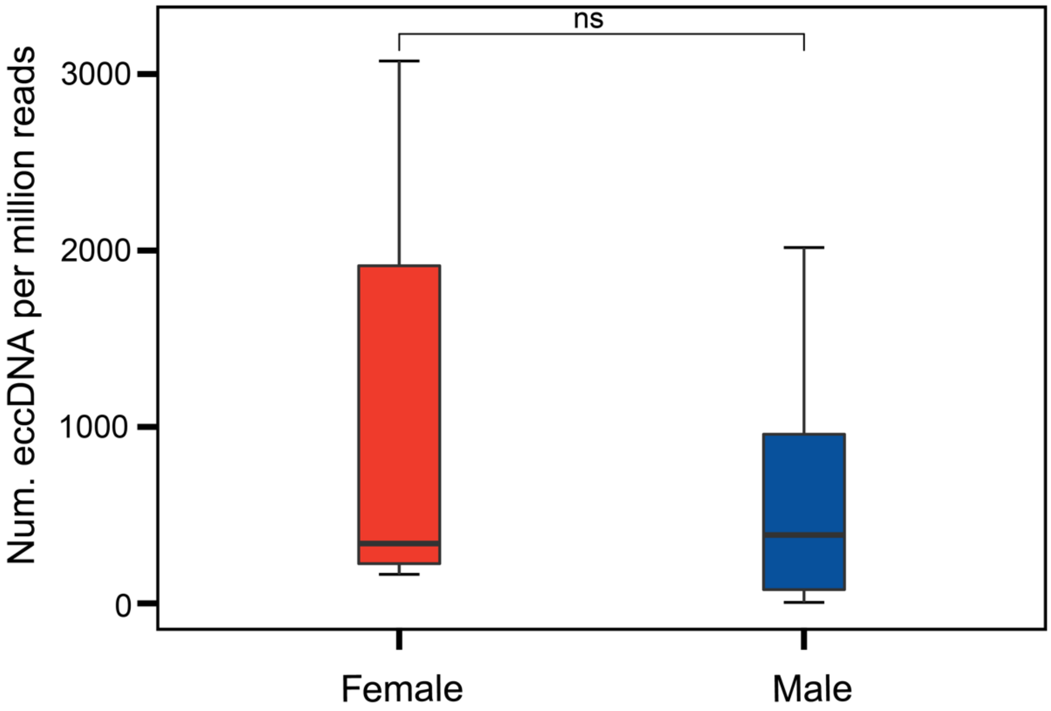


**Fig. S2.** Comparison of the number of urinary cell free eccDNA per million mapped reads between all of the men and women (P = 0.29). ns, not significant.

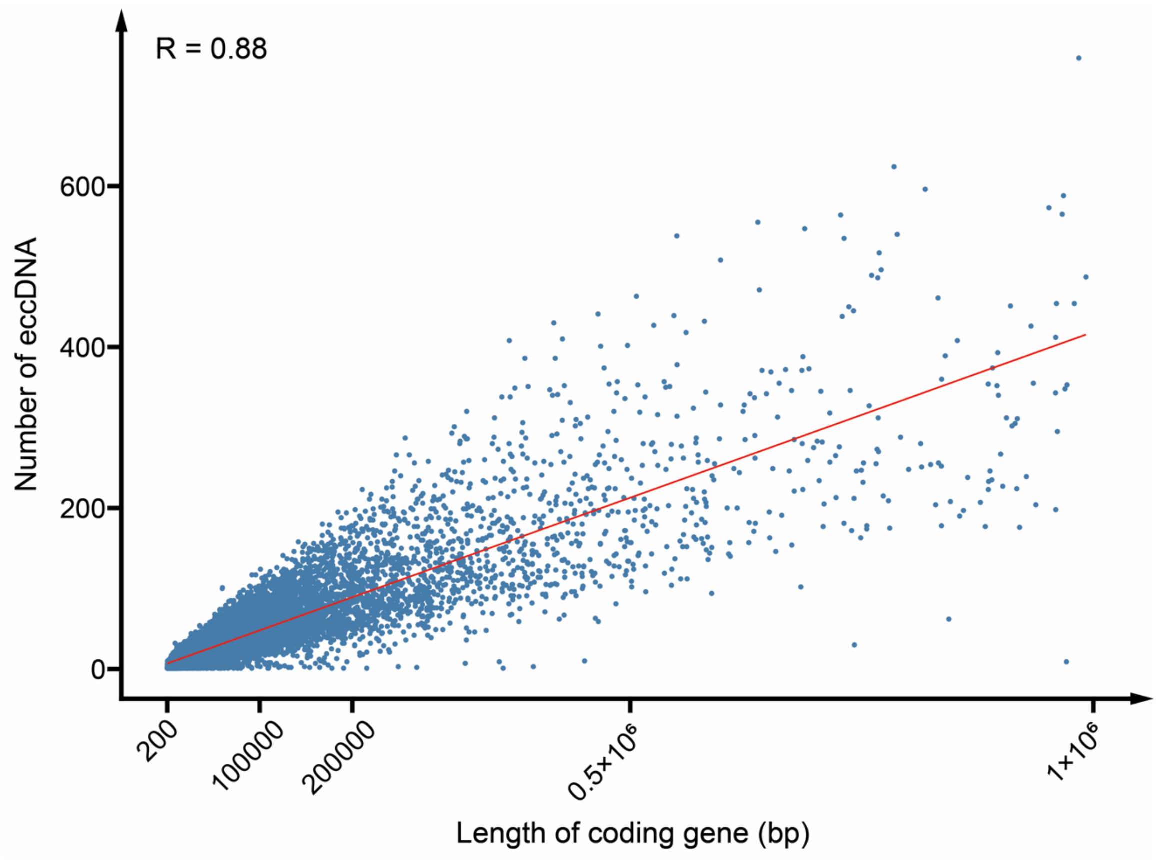


**Fig. S3.** Analysis of the correlation between the length of coding gene and the number of eccDNA.

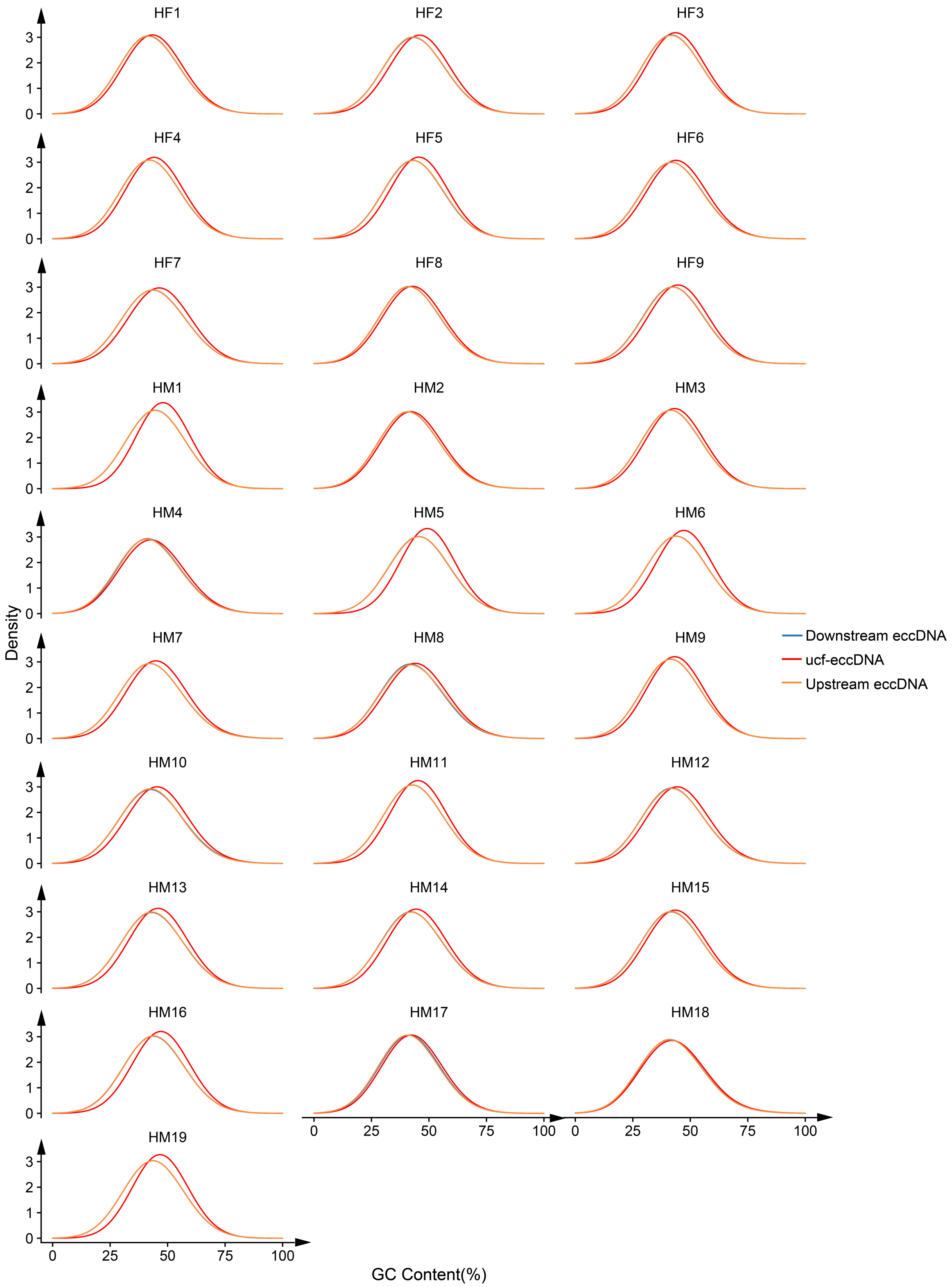


**Fig. S4.** GC content distribution of urinary cell free eccDNAs from 28 individual cases. HM: healthy male; HF: healthy female.

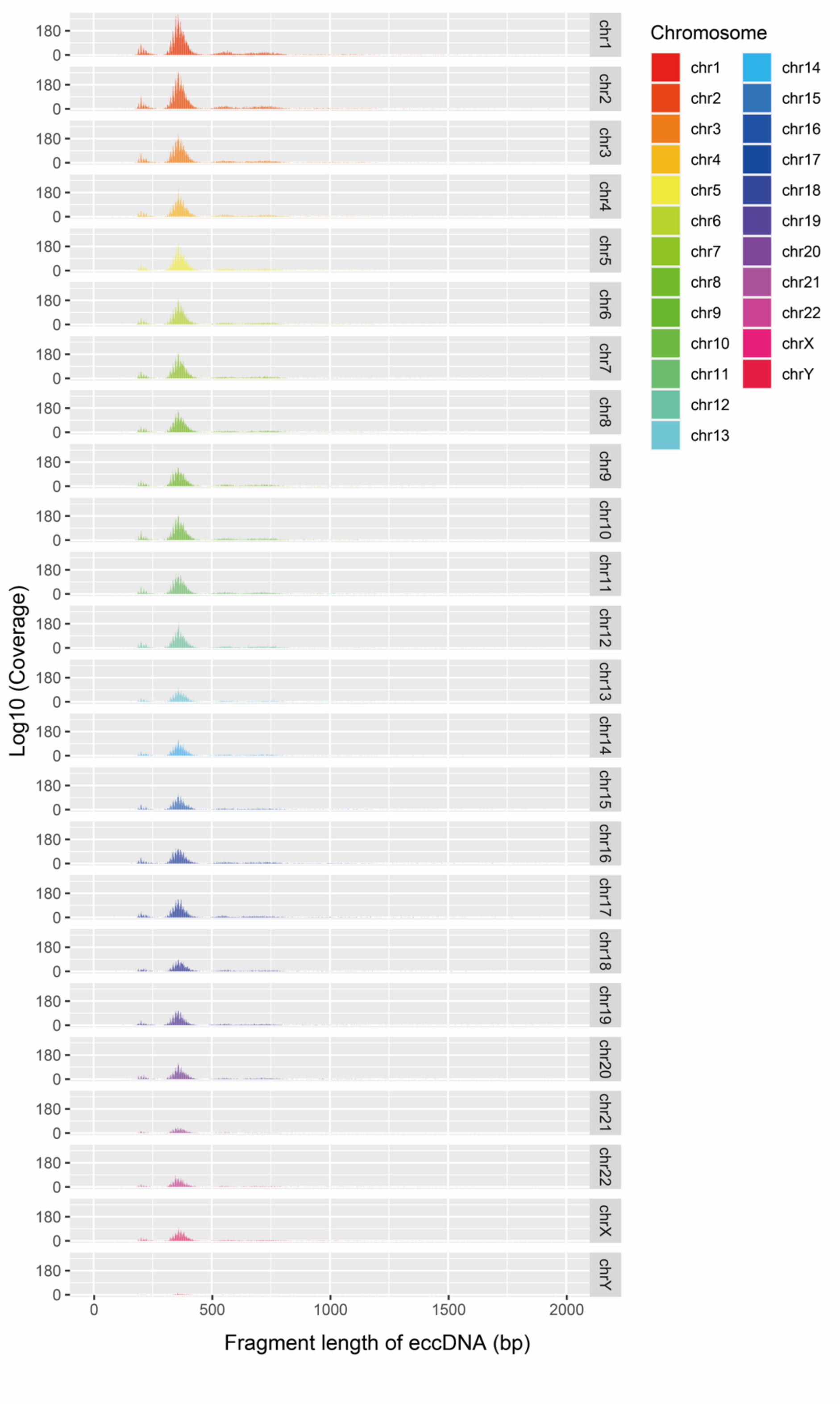


**Fig. S5.** Fragment length distribution of urinary cell free eccDNAs in each chromosome (combined data from 28 cases).

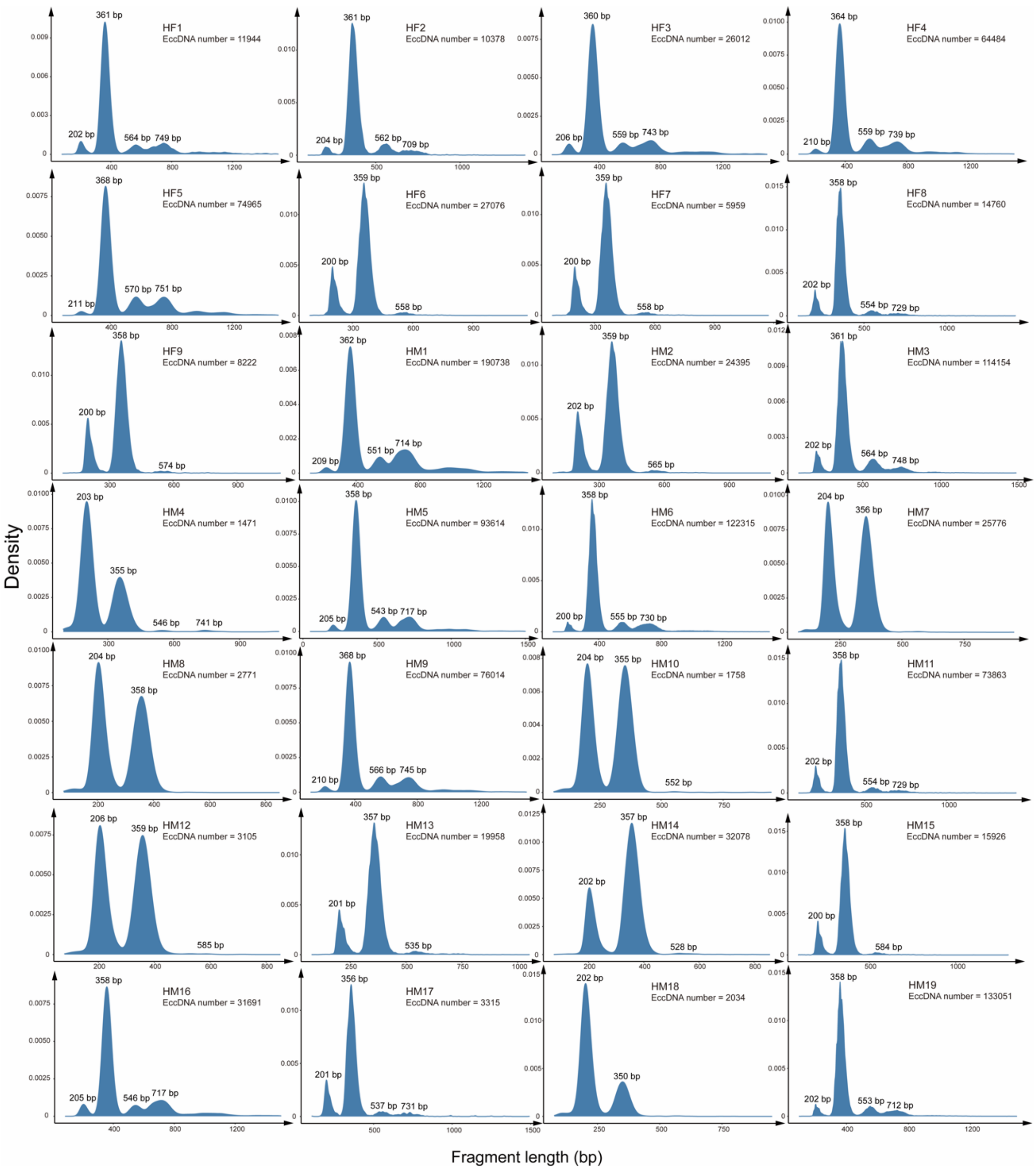


**Fig. S6.** Fragment length distribution of urinary cell free eccDNAs from 28 individual cases. HM: healthy male; HF: healthy female.

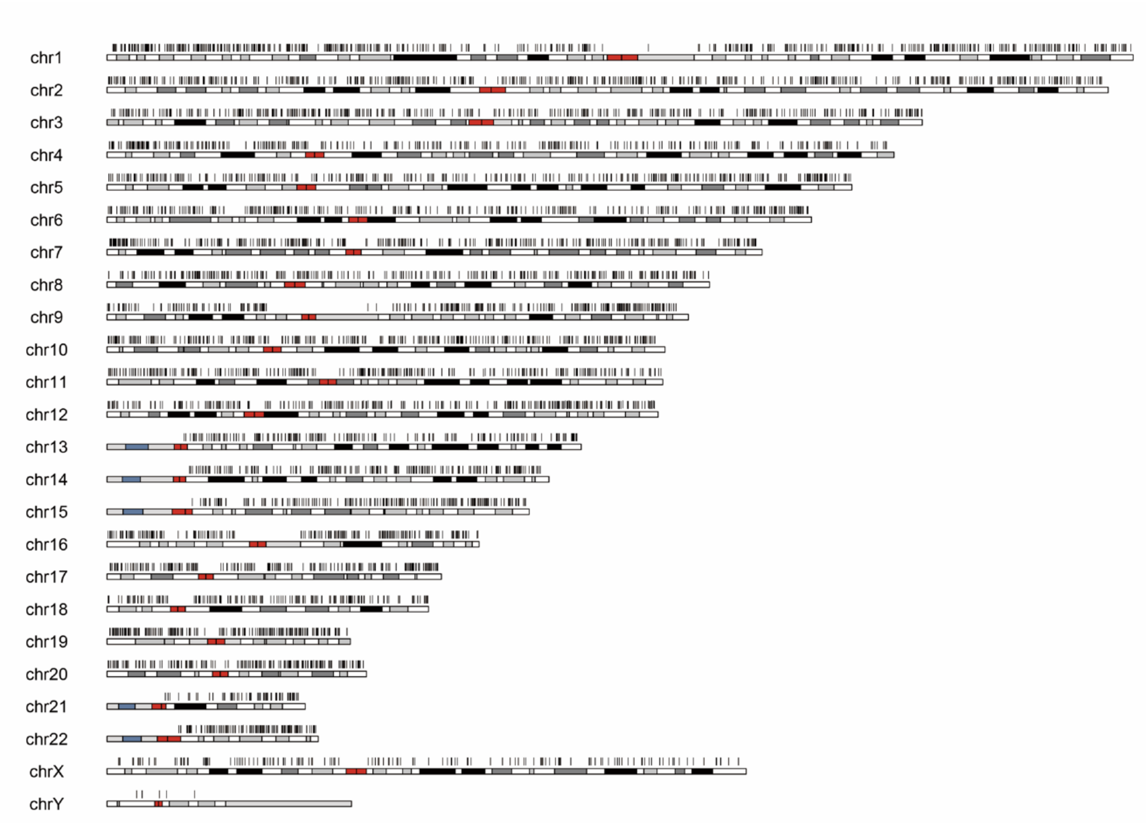


**Fig. S7.** Karyotype plot showing chromosomal distribution of randomly selected 5000 urinary cell free eccDNAs.

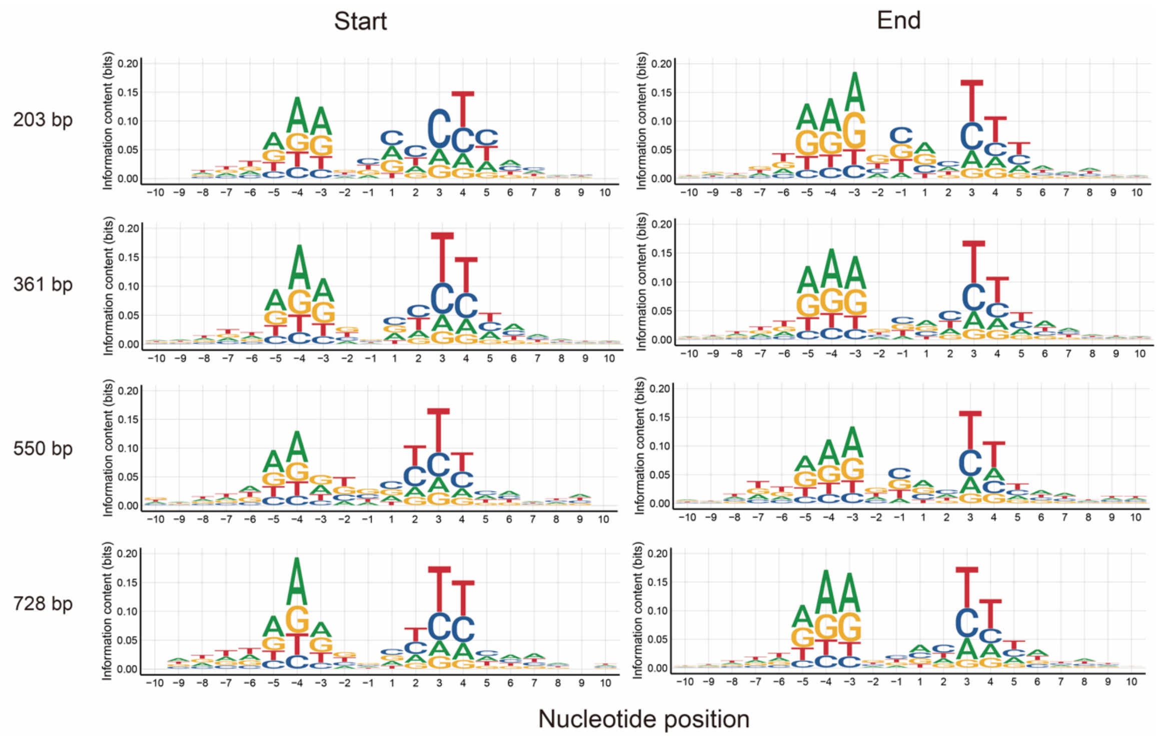


**Fig. S8.** The nucleotide frequencies surrounding the start and end sites of urinary cell free eccDNAs with different peak sizes.

| Sample | Gender | Age | number of eccDNA |
|---|---|---|---|
| HF1 | Female | 22 | 11944 |
| HF2 | Female | 26 | 10378 |
| HF3 | Female | 24 | 26012 |
| HF4 | Female | 26 | 64484 |
| HF5 | Female | 26 | 74965 |
| HF6 | Female | 32 | 27076 |
| HF7 | Female | 34 | 5959 |
| HF8 | Female | 35 | 14760 |
| HF9 | Female | 26 | 8222 |
| HM1 | Male | 26 | 190738 |
| HM2 | Male | 38 | 24395 |
| HM3 | Male | 39 | 114154 |
| HM4 | Male | 33 | 1471 |
| HM5 | Male | 32 | 93614 |
| HM6 | Male | 25 | 122315 |
| HM7 | Male | 29 | 25776 |
| HM8 | Male | 31 | 2771 |
| HM9 | Male | 36 | 76014 |
| HM10 | Male | 30 | 1758 |
| HM11 | Male | 27 | 73863 |
| HM12 | Male | 28 | 3105 |
| HM13 | Male | 28 | 19958 |
| HM14 | Male | 24 | 32078 |
| HM15 | Male | 30 | 15926 |
| HM16 | Male | 23 | 31691 |
| HM17 | Male | 30 | 3315 |
| HM18 | Male | 23 | 2034 |
| HM19 | Male | 25 | 133051 |

**Table S1.** The number of ucf-eccDNAs in each healthy volunteer. Ucf-eccDNA: urinary cell free eccDNA.
